# Exploring the onset and progression of prostate cancer through a multicellular agent-based model

**DOI:** 10.1101/2023.02.16.528831

**Authors:** Margot Passier, Maisa van Genderen, Anniek Zaalberg, Jeroen Kneppers, Elise Bekers, Andries M Bergman, Wilbert Zwart, Federica Eduati

**Affiliations:** Department of Biomedical Engineering, Eindhoven University of Technology, Eindhoven, the Netherlands; Division of Oncogenomics, Oncode Institute, The Netherlands Cancer Institute, Plesmanlaan 121, 1066 CX Amsterdam, The Netherlands; Division of Pathology, The Netherlands Cancer Institute, Amsterdam, 1066 CX, The Netherlands; Division of Medical Oncology, Netherlands Cancer Institute, Plesmanlaan 121, 1066 CX, Amsterdam, the Netherlands; Institute for Complex Molecular Systems, Eindhoven University of Technology, PO Box 513, 5600MB, Eindhoven, The Netherlands

**Keywords:** Prostate cancer, agent-based modeling, cancer development, tumor microenvironment, multicellular model

## Abstract

Over ten percent of men will be diagnosed with prostate cancer (PCa) during their lifetime. Arising from luminal cells of the prostatic acinus, PCa is influenced by multiple cells in its microenvironment. To expand our knowledge and explore means to prevent and treat the disease, it is important to understand what drives the onset and early stages of PCa. In this study, we developed an agent-based model of a prostatic acinus including its microenvironment, to allow for *in silico* studying of PCa development.

The model was based on prior reports and in-house data of tumor cells co-cultured with Cancer Associated Fibroblasts (CAFs) and pro-tumor and/or anti-tumor macrophages. Growth patterns depicted by the model were pathologically validated on H&E slide images of human PCa specimens. We identified that stochasticity of interactions between macrophages and tumor cells at early stages strongly affect tumor development. Additionally, we discovered that more systematic deviations in tumor development result from a combinatorial effect of the probability of acquiring mutations and the tumor-promoting abilities of CAFs and macrophages. *In silico* modeled tumors were then compared with 494 cancer patients with matching characteristics, showing strong association between predicted tumor load and patients’ clinical outcome. Our findings suggest that the likelihood of tumor formation depends on a combination of stochastic events and systematic characteristics. While stochasticity cannot be controlled, information on systematic effects may aid the development of prevention strategies tailored to the molecular characteristics of an individual patient.

## Introduction

Prostate cancer (PCa) is generally diagnosed at late age, with 75% of all cases found in men over 65 years old (1,2), while the formation of precursor neoplastic lesions is initiated years earlier (3). While localized PCa can be cured, metastatic disease cannot, and its treatment is a clinical challenge (4,5). Currently, PCa is the second most diagnosed cancer and the second leading cause of cancer deaths in men globally (1). Studying the onset and early development of PCa improves our understanding of this disease and could aid the development of new treatment strategies to prevent disease progression and to improve clinical care for PCa patients (6–10).

PCa generally initiates in the prostatic acini. In a normal acinus the epithelium is highly organized with a bilayer of basal and luminal cells separated from the underlying stroma by the basement membrane. During the premalignant prostatic intraepithelial neoplasia (PIN) stage, luminal cells start to hyperproliferate (11,12). Eventually, this can lead to the disruption of the basal cell layer and breakdown of the basement membrane, which is a prerequisite for the invasion of tumor cells into the tumor microenvironment (TME) (13,14), allowing cancer cells to metastasize (15,16).

PCa is assumed to originate from mutations that confer a proliferative advantage to the transformed cells (17,18). The accumulation of mutations is essential for the progression towards the malignant disease, and PCa is characterized by a high heterogeneity of tumor cells (19,20), with clonal selection shaping tumor evolution (21). Fibroblasts, normally contribute to maintenance of the healthy homeostasis in the prostate (22–24). However, when in contact with neoplastic cells they can differentiate into cancer-associated-fibroblasts (CAFs) (22). CAF differentiation already occurs in early premalignant stages, contributing to the development and progression of PCa by stimulating tumor cell proliferation (25) and migration (26,27) and by altering the surrounding extracellular matrix (28–30), facilitating cancer cells to invade the stroma (31,32). Macrophages are another important cell type in PCa development, constituting 70% of the immune cell population in the prostate TME (33). Macrophages are attracted by cytokines released by PCa cells and initially contribute to the immune defense against tumors (34). However, macrophages have a wide range of functions depending on environmental cues and can differentiate from a pro-inflammatory and anti-cancer (M1-type) to a pro-cancer (M2-type) phenotype (35). The latter may support tumor cell proliferation, migration, and invasion (36,37).

Although several studies have characterized developmental stages of PCa and the underlying molecular mechanisms of tumorigenesis (12,18,35,38–40), it is still unclear how such mechanisms jointly contribute to PCa development (41).

Given the limitations of *in vivo* temporal data acquisition in studying heterogeneity at early stages in patients, novel models are required to study development of PCa. Mathematical models offer valuable tools to study tumor development *in silico*. In particular, agent-based models (ABM) are spatial models that simulate the effect of interactions in complex multicellular systems such as tumors. This enables the investigation of how the overall system behavior originates from the interaction of individual components (42). In ABMs, cells are seen as agents that can interact with the surrounding cells (agents) based on a predefined set of rules. Based on stochastic simulations, ABMs enable monitoring the evolution of the tumor over time, and systematically test the impact of different aspects of the TME in a controlled way that would be unfeasible in any *in vitro* or *in vivo* settings (43).

Here we propose the first comprehensive ABM of PCa onset and progression encompassing nine agent types and 60 parameters. Our model parameters are based on prior reports and in-house generated experimental data on LNCaP cultures and cocultures with fibroblasts, pro-tumor, and anti-tumor macrophages. We show that our model reliably recapitulates different stages and spatial morphologies observed in cancer development, based on strong phenotypical parallels with histopathology images from PCa patients. Additionally, we use the model to study which factors in the microenvironment mostly affect PCa development, and to simulate *in silico* patients with different molecular characteristics, showing strong associations between *in silico* tumors and matching clinical data from The Cancer Genome Atlas (TCGA). We provide our ABM as a tool to systematically study the impact of the microenvironment on PCa development.

## Materials and Methods

### Agent-based modeling assumptions and simulations

In this study we developed two ABMs to: 1. Test the requirements for PCa tumor maintenance and 2. Study the onset and progression of PCa. In both cases we used a two-dimensional (2D), on grid, stochastic ABM. The size of one grid space was set to the size of one tumor cell, 142.89 *μ*m2 (44) forming a 125×125 grid. The first model only includes tumor cells (normal and stem-like) and in all scenarios a total of 1500 cells were randomly seeded. The second model includes nine different types of cellular agents (i.e., different *in silico* cell types) and cells were no longer seeded randomly, but in an ellipsoid geometry, mimicking the prostatic acinus. The average size of the lumen of the acinus was determined at 73 μm (6 gridspaces) (45) and increased to 156 μm (13 gridspaces), to adapt for the limitation that there are only two directions for growth and migration in the 2D model. Simulations were always repeated multiple times (as specified in the corresponding results sections) to account for the stochastic nature of ABM simulations.

Like all models, our models are an abstraction of reality and based on a set of assumptions which are listed in **Supplementary Table 1**. All agents (cells) occupy one space on the grid and compete for space in their Moore neighborhood (i.e., the eight surrounding grid spaces). The model iterates through a defined number of time steps. At each step every agent can perform an action with a certain predefined probability. These probabilities are defined by model parameters which are either derived from literature or estimated from our experimental data as detailed in the next sections. The complete list of model parameters is provided in **Supplementary Table 2**.

### Modeling of tumor cells as cellular agents

In both models, tumor cell agents are seen as mutated luminal cells (normal or stem cells) and they have the possibility to acquire mutations (probability defined by the model parameter *TUpmut*; **Supplementary Table 2**) which confers them a proliferative advantage modeled as a (cumulative) increase in the probability of proliferation and maximum proliferation capacity (*TUadded values*) (17). Mutated cells can migrate (*TUpmig*), die (*TUpdeath*) or proliferate (*TUpprol*). Cancer stem cells have the same characteristics as normal tumor cells, but they are additionally characterized by their self-renewal capacity (46). Therefore, stem cells are modeled as having infinite proliferation capacity, while other luminal cells have a limited proliferation capacity (*TUpmax*).

### Implementation of an agent-based model of PCa onset and progression

The more complex ABM that we developed to study PCa developments includes the tumor cells described in the previous section, and eight additional agents that can perform actions and interact with each other (**Fig. 1**). As stated above, this model’s starting geometry mimics the one of a healthy prostate acinus, where luminal cells (including a fraction of stem cells) are placed on a layer of basal cells, which is attached to the basement membrane (47,48). Luminal cells can acquire mutations and convert into tumor cells. A layer of tissue resident fibroblasts is placed outside of the acinus, surrounded by extracellular matrix (ECM) containing more fibroblasts. Fibroblasts can convert to tumor-promoting CAFs when they are in proximity of tumor cells (22,39,49,50). Macrophages can enter the simulation from the top left corner, simulating entry from a blood vessel. Although they exist in a broad spectrum, we consider a simplification of two phenotypes: M1 (immuno-promoting/anti-tumor) and M2 (tumor-promoting, or TAMs) macrophages (51).

**Figure 1.**
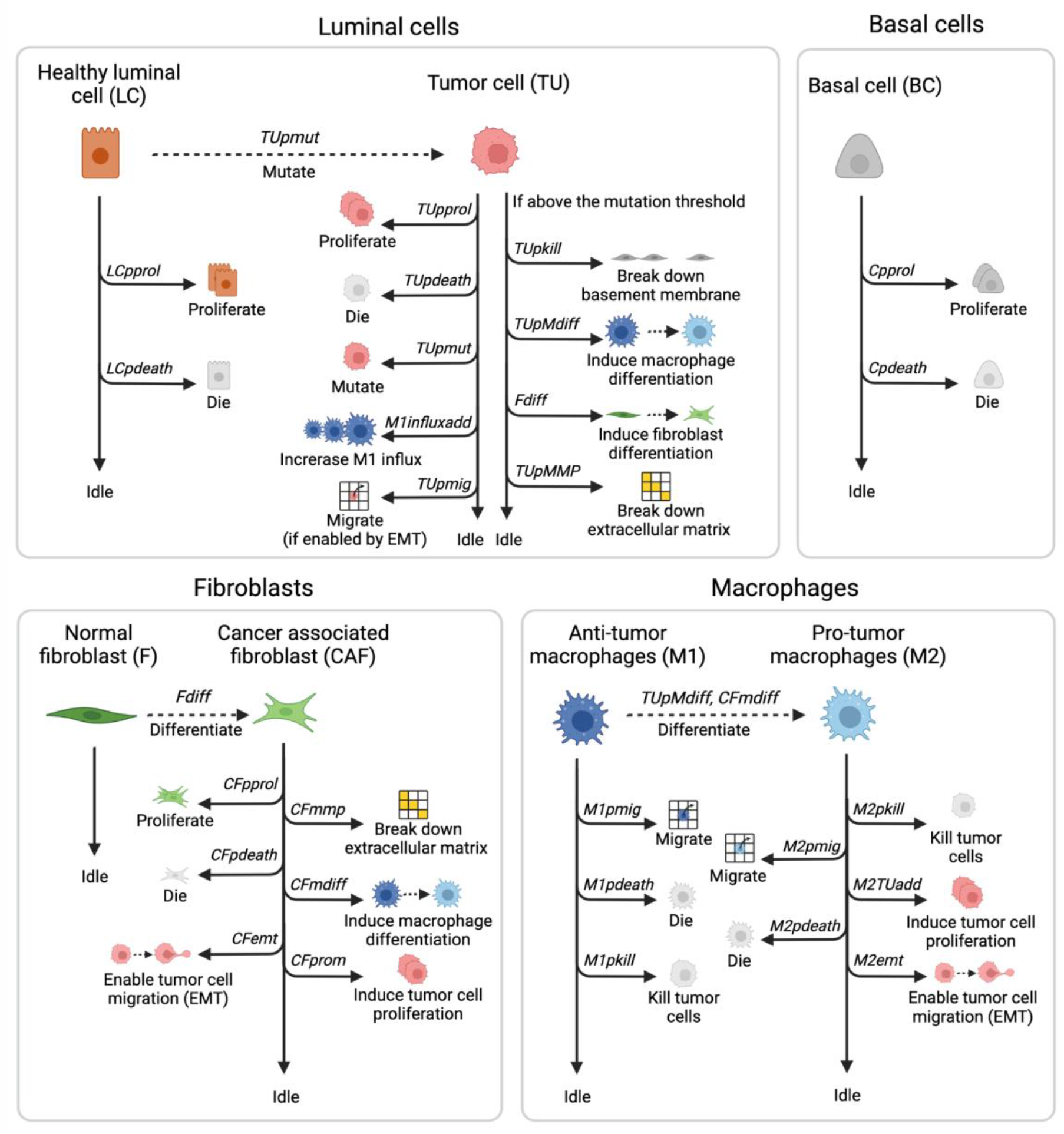
Overview of the agents and actions they can perform during each model iteration. The simulation starts with luminal cells (LC) and basal cells (BC) that can proliferate, die, or idle, all within physiological regions and with fixed probabilities. The starting geometry also contains quiescent fibroblasts (F) and the passive agents (basement membrane and ECM), macrophages enter throughout the simulation. LCs can gain mutations, resulting in an increased M1-macrophage influx, once sensed by macrophages. These mutated cells (TU) can additionally break down basement membrane and ECM and affect macrophage and fibroblast differentiation upon reaching mutation thresholds. Differentiated fibroblasts (CAF) proliferate, die, and can perform tumor-promoting actions. Just as the differentiated M2 macrophages, they stimulate TU proliferation and allow for TU migration. Macrophages (M1 and M2) can also kill tumor cells and die or migrate. Image created with BioRender.com.

In each iteration, all agents have their own round during which they can perform their actions or can idle based on the defined probabilities. The basement membrane and the ECM are instead passive agents that can only idle or be affected by the actions of other agents. Actions are performed by agents in the following order.

- Luminal cells can proliferate (*LCpprol*) within their physiological region and die (*LCpdeath*). They can also gain mutations (*TUpmut*), thereby converting into tumor cells (47). Tumor cells can die (*TUpdeath)*, proliferate (*TUprol)* also outside their physiological region, affect fibroblast differentiation (*Fdiff*) and increase macrophage influx (*M1influxadd*) (37,52). Additionally, they can gain more mutations (*TUpmut)*. Upon reaching mutation thresholds (*TUthrshBM, TUthrshM, TUthrshMMP*), tumor cells can perform additional actions: break down the basement membrane (*TUpkill)*, affect macrophage differentiation (*TUpMdiff)*, or break down the ECM (*TUpMMP) (48,51,53)*. After going through epithelial-mesenchymal transition (EMT), which is promoted by CAF or TAM proximity, tumor cells become invasive and can migrate randomly to an empty space in the Moore neighborhood (*TUpmig*) (37,54,55).
- Basal cells can proliferate within their physiological regions (*Cprol)* and die (*Cpdeath)*. They must remain attached to the basement membrane to survive and cannot invade the lumen (56).
- Fibroblasts are quiescent, i.e., they only idle (57). However, when they are in close proximity to tumor cells (i.e. max two grid spaces away, so the tumor cells can affect fibroblast differentiation over the basement membrane during PIN), they can differentiate into CAFs (*Fdiff*) (58). CAFs can proliferate (*CFpprol)*, die (*CFpdeath)*, break down ECM (*CFmmp)*, promote differentiation of macrophages towards the tumor-promoting phenotype (*CFmdiff)*, enable migration for mutated cells (*CFemt)* and promote tumor cell proliferation (*CFprom)*, by adding to the proliferation probability of tumor cells (25,53,54,58).
- Macrophages can enter the simulation (*M1influxProb*), with an increased probability when macrophages detect tumor cells (*M1influxadd*) (37,52). All macrophages enter the simulation as M1 macrophages that can kill tumor cells (*M1pkill)*, die (*M1pdeath)* or migrate (*M1pmig)*. Macrophages move randomly, unless they can sense (within 17 grid spaces, to account for the effect of chemokines) tumor cells, as they will then move towards them (59,60). When differentiated into tumor-promoting M2 macrophages, via stimulation by tumor cells or CAFs, they can additionally promote tumor cell proliferation (*M2TUadd)* and enable tumor cell migration (*M2emt) (37)*.

For typical simulations in this study, steps of 12 hours were used to simulate a period of 400 days. At each step the model iterates through the rounds described above and each agent can perform one or more actions. Apart from stem cells, all other cells have a maximum number of times they can proliferate (luminal cells, tumor cells, basal cells, fibroblasts and CAFs) or kill (macrophages) after which they get exhausted and die. Migration and proliferation can only occur in the standard Moore neighborhood, except for macrophages that can migrate in the Moore neighborhood of range two (24 neighbors instead of 8), to allow for acinus infiltration by traveling over the basement membrane (37,52,60).

### Experimental data for parameter estimation

We performed co-culture *in vitro* experiments for fitting the model parameters. We used the PCa cell line LNCaP (ATCC), immortalized foreskin fibroblast cells (BJ fibroblasts, Agami Lab NKI) and the monocytic cell line THP-1 (ATCC) which were differentiated into M1 or M2 macrophages.

LNCaP cells and fibroblasts were cultured together with either M1- or M2-macrophages in a 4:1:1 ratio respectively. Cells were cultured in physiological hormonal conditions with R1881 used to induce androgen receptor (AR) signaling. LNCaP cells were tagged with eGFP to follow them overtime. LNCaP-eGFP cell proliferation was measured with IncuCyte Zoom fluorescent signal imaging system for seven days and performed in triplicate. Lastly, BJ fibroblast proliferation was measured separately by analysis of phase-contrast images from IncuCyte Zoom to obtain fibroblast growth curves, for fibroblast parameter determination.

Apoptosis was measured in real time using IncuCyte Zoom (Essen, BioScience). To this end, cells were grown in FBS, including androgen, with an addition of Caspase-3/7 Read Reagent for Apoptosis (Essen Bioscience) in duplicate.

The resulting growth curves (**Supplementary Fig. S1**) and apoptosis data of PCa cells were used to determine the parameters of tumor cells in the model.

### Parameter identification

Tumor cell, fibroblast and macrophage parameters were estimated using particle swarm optimization (PSO) to fit the experimental data (**Supplementary Fig. S1**). For each biological replicate, parameters were optimized 50 times to account for biological variation and model stochasticity. Final parameter values were fixed to the average estimated value after assessing the robustness of the estimated values between replicates. The optimizations were done sequentially, fixing the estimated model parameters. First, *TUpmax* and *TUpprol* were fitted to the experimental growth curves of the LNCaP cells. *TUpdeath* was determined by measuring apoptosis of LNCaP cells. Subsequently, *Fpprol, Fpmax* and *Fpdeath* were fitted using the fibroblast growth curves. Lastly, *M1pkill* and *M1kmax* were fitted using the experimentally obtained growth curve for tumor cells in the presence of M1 macrophages and fibroblasts. Similarly, *M2pkill was* determined. *M2kmax* was assumed equivalent to *M1kmax*.

The remaining parameter values were either derived from previous studies, adapted from a previously published model of colorectal cancer (60,61) or qualitatively tuned (all details and specific references are in **Supplementary Table 2**).

### Parameter sensitivity analysis

A qualitative sensitivity analysis was performed for all individual model parameters by increasing them individually by 10% and recording the percentage change in output, in the number of tumor cells at 400 days. All simulations were conducted ten times to account for model stochasticity. Parameters with low sensitivity (i.e., for which the increase did not affect the output above the deviations due to the stochasticity of the model) were fixed and are specifically mentioned in **Supplementary Table 2**. Follow up analysis were conducted for the four most sensitive parameters (i.e. those causing on average > 10% change in output), simulating ten intermediate values in the region of interest (i.e., in which the effect of changing the parameter is visible but not so extreme as to overpower all other parameters). Lastly, pairwise combinations (with five parameter values each) of the most sensitive parameters were conducted to see if there were synergistic or antagonistic relations. In all sensitivity analyses the relative tumor size was recorded at 400 days and averaged across ten simulations.

### Pathology slides for assessment of morphological features

Pathology slices of PCa patients were used, with permission, to compare growth patterns in patients with the model simulations. The patient samples were randomly picked out of daily practice of prostatectomies of PCa patients. Every slide consists of a 4 μm thick section of FFPE material and was stained with haematotoxylin and eosin (H&E). The uropathologist scanned the slides and chose representative images of prostate carcinoma.

### Comparison between model simulations and clinical patient data

Model predictions were compared with clinical data from The Cancer Genome Atlas (TCGA). We used a cohort of N=494 PCa patients for which molecular data (transcriptomics and genomics) and survival data (62) were available. RNA sequencing (RNA-seq) data was downloaded via the Firehose tool from the BROAD institute (released January 28, 2016) and processed as described by Lapuente-Santana et al (63). To allow for comparison between expression levels of different genes, transcripts per million (TPM) were used. Tumor mutational burden (TMB) data were obtained from a previous report (64). Quantifications of the relevant cell types for individual patients were obtained using deconvolution methods accessible through the *immunedeconv* R package (65): M1 and M2 macrophages were obtained using quanTIseq (66) and CAFs were derived using EPIC (67). Lastly, for 333 PCa patients we also retrieved information on Gleason score and binarized them as high (>=7) and low (<7) Gleason score (68). For the comparison of model simulations and clinical Progression Free Survival (PFS) we used correlation analysis (Spearman and Pearson) and Kaplan Meier plots (using *survival* and *survminer* R packages).

### Computational implementation

The ABM of PCa onset and development is available as Matlab code in a GitHub repository at https://github.com/SysBioOncology/ABM_prostate_cancer_development.

## Results

### *In silico* prostate tumors require a proliferative advantage of mutated cells additionally to cancer stem cells to maintain themselves at realistic stem cell percentages

Cancer stem cells are known to play an important role in PCa development (69–73). To identify the percentage of stem cells needed for our *in silico* tumors to maintain themselves, we used a simple ABM including only normal tumor cells and/or tumor stem cells (as defined in **Material and Methods**) that were randomly seeded on the grid to test different possible scenarios *in silico* (74). For the first scenario, tumor cells were not allowed to gain a proliferative advantage via mutations. This allowed us to assess the ability of stem cells alone to sustain the tumor. Irrespective of the starting percentage of stem cells, we achieved an almost full grid at approximately 15000 tumor cells and stabilizing stem cell percentage at approximately 17% (**Fig. 2A**). While the tumor was able to survive with stem cells alone, this final stem cell percentage is much higher than we could reasonably expect based on literature, which is reported to be 0.1-0.3% in the human prostate (69). The second scenario included no stem cells, but only tumor cells with a possibility of gaining (more) mutations that confers proliferative advantage. For all simulations all tumor cells died within 40 days, meaning that a tumor cannot survive based on acquired mutations only (**Fig 2B**). The third scenario included both a percentage of initial stem cells and tumor cells with the ability of gaining mutations. In this case, the tumor could maintain itself while the percentage of stem cells stabilized at a much lower value; approximately 0.5% (**Fig. 2C**). Based on these observations, we conclude that the combination of stem cells and possibility for luminal cells to mutate (and with that, gain a proliferative advantage), is required for tumor maintenance at realistic stem cell levels, and that this does not depend on the initial percentage of stem cells.

**Figure 2.**
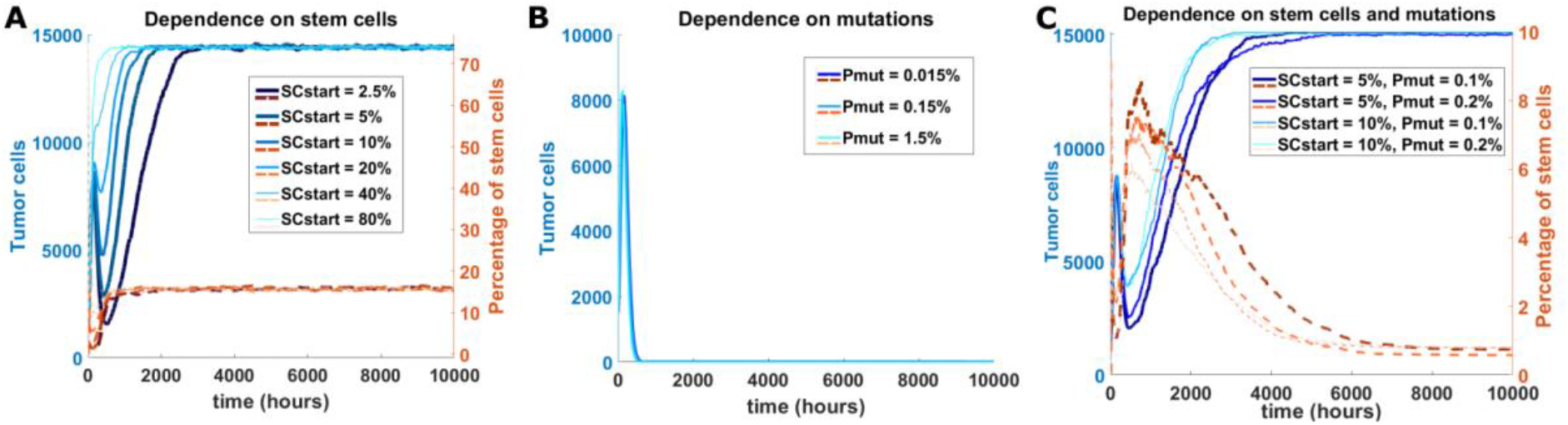
In silico testing of requirements for tumor maintenance. **A**, Amount of tumor cells (blue) and percentage of stem cells (orange, dotted) simulated over time under the condition that included only stem cells to maintain tumors. Simulations for six different initial percentages of stem cells (SCstart) are shown. **B**, Similar plot testing the condition in which the proliferative advantage of mutated tumor cells is the only source for tumor maintenance. Simulations for three different probabilities of acquiring mutations (Pmut) are shown. **C**, Similar plot testing the condition in which tumor maintenance depends on both stem cells and tumor cells that can gain mutations. Four combinations of initial stem cell percentage and probability of mutation acquisition are shown.

### Model simulations recapitulate known steps of PCa development

After defining the basic requirements for tumor maintenance, we developed a comprehensive ABM to describe onset and development of PCa in a simulated *in vivo* setting starting from a healthy prostate acinus (**Fig. 3**). This model is schematically depicted in **Fig. 1** and is based on the set of assumptions and parameters in **Supplementary Table 1** and **2** respectively (see **Material and Methods**).

**Figure 3.**
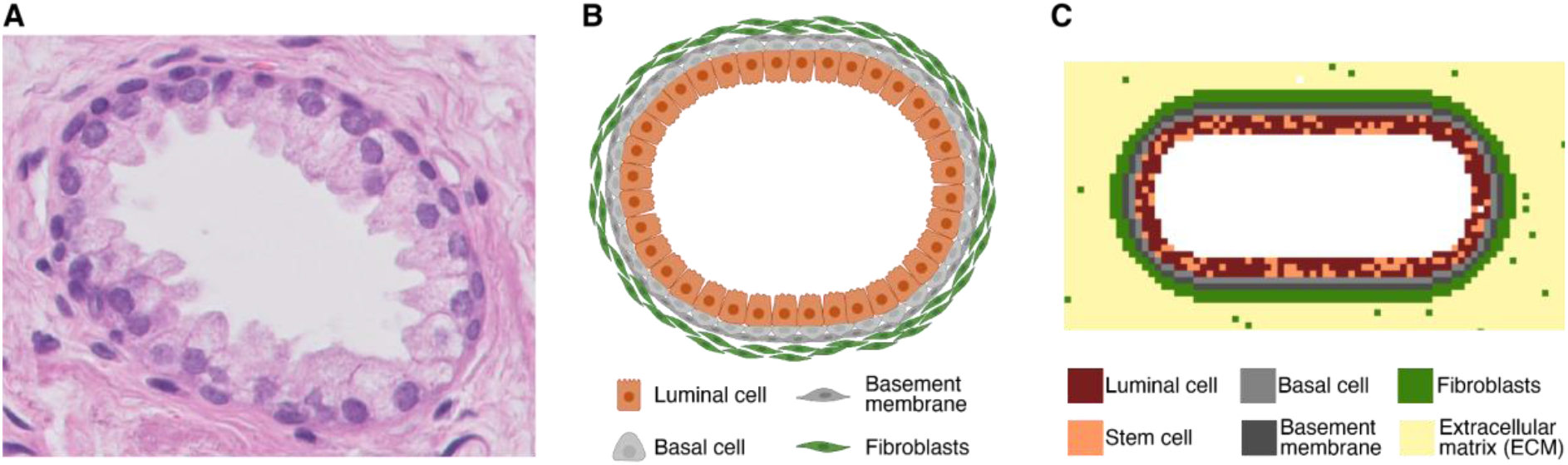
Overview of the starting geometry in threefold; a pathology slice, schematic representation, and model geometry visualization. **A**, A histology slice of a healthy prostatic acinus (H&E staining, 400x magnification). **B**, Schematic representation of the acinus. **C**, Modeled starting geometry, including a color scheme of all cells included in the starting geometry

Running the model simulations, we can observe how PCa develops over time (**Fig. 4A-I**, video in **Supplementary File V1)**. The initial condition is a healthy prostatic acinus with empty lumen **(Fig. 4A)**. Luminal cells can start to mutate and then grow in the lumen (**Fig. 4B**). Mutated luminal cells give rise to prostatic intraepithelial neoplasia (PIN), characterized by luminal cell hyperplasia, while the basement membrane remains intact (19,75,76) (**Fig. 4B-F**). Mutated luminal cells (hereafter called tumor cells) attract macrophages, resulting in an increased macrophage influx towards the acinus (**Fig. 4C**) (37,51). Basal cell layer breakdown starts to occur during early PIN (**Fig. 4D**) and increases exponentially with disease progression (16). During PCa development, CAFs originate from normal fibroblasts due to tumor cell stimulation (**Fig. 4E**) (77). Tumor cells also affect polarization of macrophages towards the tumor-promoting phenotype by cytokine secretion, resulting in an increased number of M2-like macrophages (**Fig. 4F**). This increasing tumor-promoting environment results in basement membrane breakdown (**Fig. 4G**) allowing the disease to progress towards cancer. The tumor promoting cells (TAMs and CAFs) elicit EMT in tumor cells, making them invasive (**Fig. 4H**) (54,55). This results in tumor cells invading the surrounding tissue, and thereby starting the cancerous phase (**Fig. 4I**). Based on these findings, we conclude that our model can represent all main steps of PCa onset and development well.

**Figure 4.**
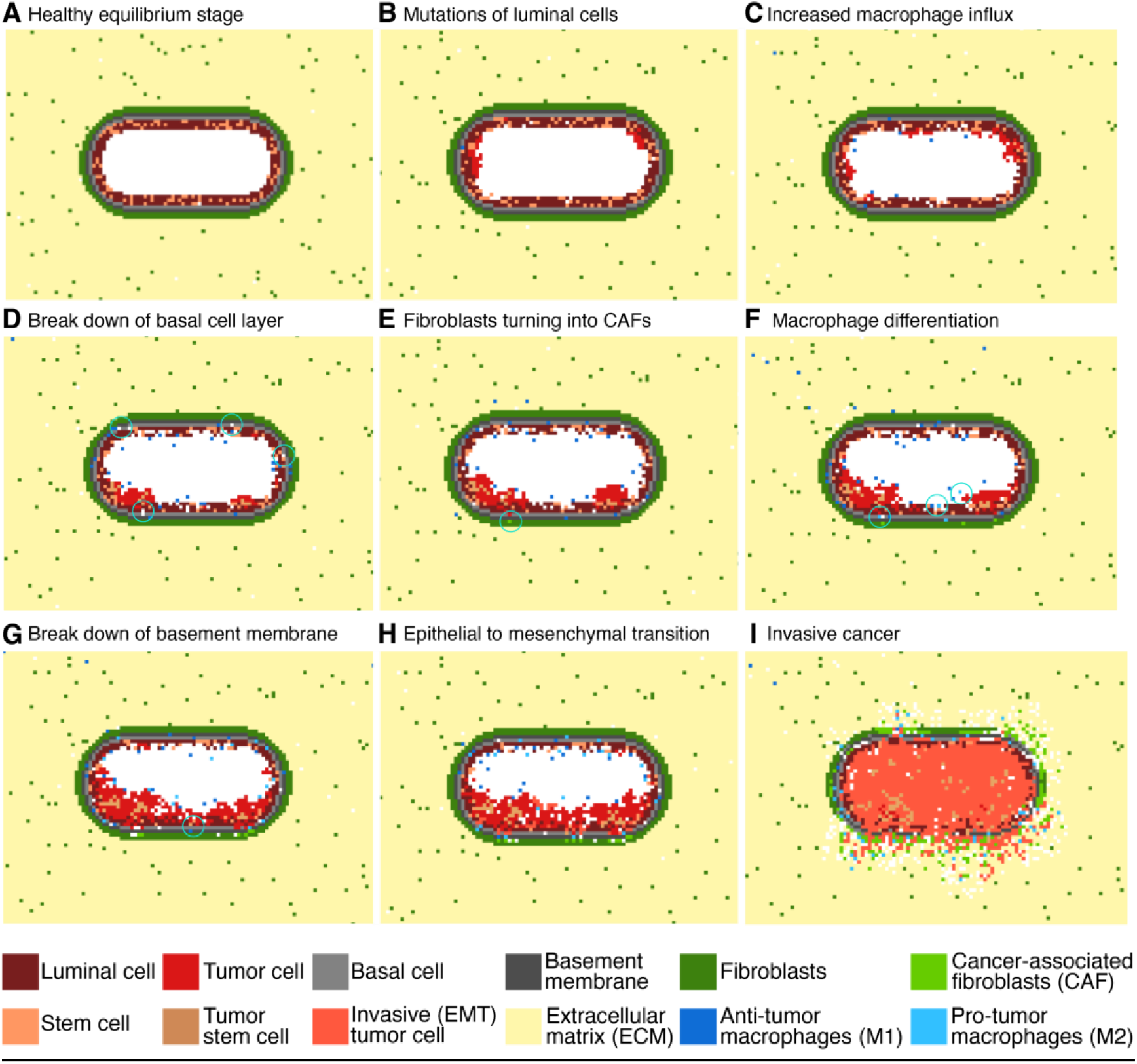
Initial healthy stage and following eight steps of PCa development as by PCa ABM simulation. **A**, Healthy prostatic acinus. **B**, Mutations start to occur in the luminal cells converting them into tumor cells. **C**, The presence of mutated cells increases the influx of M1 macrophages. **D**, Mutated cells start to occupy spaces in the basal cell layer. **E**, Fibroblasts are differentiating towards their tumor-promoting phenotype (CAFs). **F**, Macrophages are differentiating towards their tumor-promoting phenotype. **G**, All these factors lead to break down of the basement membrane. **H**, Mutated cells become more invasive and start undergoing EMT. **I**, Invasive cancer with cells spreading through the surrounding tissue. The white grid spaces indicate ‘empty space’, corresponding to the lumen or to the cleaved ECM (for example by CAFs).

Using the parameter set defined in **Supplementary Table 2**, we ran 500 simulations and observed that only 36% of them results in breaking down of the basement membrane, which we consider as a marker of invasive PCa. We decided to investigate the main stochastic factors contributing to tumor development *in silico*. If the malignant cells are recognized by the macrophages at an early stage, this results in a fast increase in the ratio of macrophages to tumor cells. This allows the immune system to control and overcome the disease (**Supplementary Fig. S2A**). However, if this does not happen at early stages, the tumor develops to evade the immune response and subverts the immune response by converting macrophages to the pro-tumor phenotype, increasing the M2:M1 macrophage ratio (**Supplementary Fig. S2B**). We also observed that there are several factors that contribute to determining the time of invasion. Earlier invasions are characterized by higher numbers of CAFs, a higher average mutation load and higher M2:M1 macrophage ratio (**Supplementary Fig. S2C-E**). These results highlight how, based on stochastic simulations, our ABM enabled us to identify the aleatory factors that support PCa development.

### Model simulations recapitulate geometries present in histology images

Does our *in silico* prostate cancer model reliably represent clinically observed tumor growth patterns? To address this question, we compared our model simulations with pathology slides of PCa patients that were randomly picked out of daily practice. The uropathologist scanned the slides and selected representative images of prostate carcinoma. A common growth pattern during the PIN phase is tufting, which is characterized by protrusions consisting of multiple cell layers growing on the basal cell layer (78) (**Fig. 5A**), which was observed as emergent behavior in our model simulations (**Fig. 5B**). In the simulations, this tufted geometry originates from mutated cells that grow in clusters attached to the basal cell layer. Interestingly, permanent ‘tufts’ in our model contain stem cells suggesting that the presence of stem cell clusters could be an indication of the directionality of tumor growth. Another common growth pattern in developing PCa is bridging, when cells grow from one side of the acinus towards the other side (**Fig. 5C**), which was also portrayed in the *in silico* developing tumors (**Fig. 5D**). Overall, we can conclude that our ABM recapitulates important growth patterns observed in histology slices of actual PCa patients.

**Figure 5.**
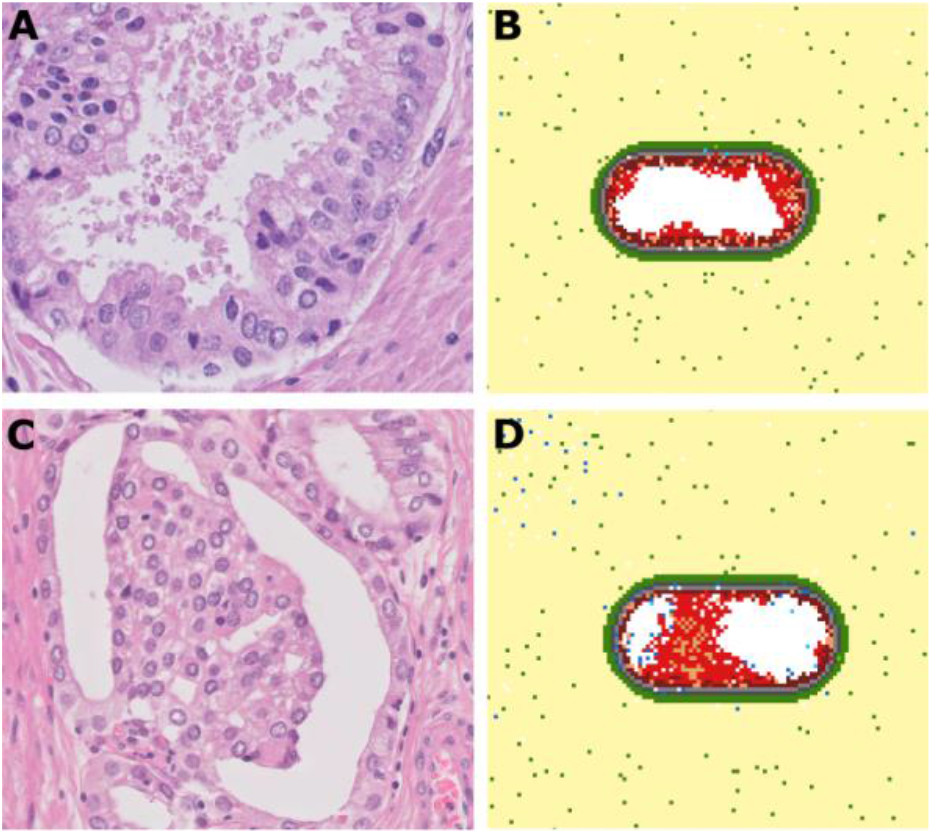
Comparison between model simulations and histology images (tufting and bridging). **A**, Pathology slice of a PCa patient (H&E staining, 400x magnification) showing a ‘tufted’ pattern of growths on the luminal cell layer. **B**, Model simulation depicting the tufting growth pattern. **C**, Pathology slice of a PCa patient (H&E staining, 400x magnification) showing bridging; growth of cells from one side of the acinus towards the other side. **D**, Simulated PCa development showing the bridging growth pattern.

### Tumor development is most strongly impacted by mutation probability, tumor promoting ability of CAFs and macrophage phenotype

Having established that the simulated onset and development of PCa recapitulates tumor developmental processes and growth patterns as observed in patients, we next investigated which model parameters most strongly affect tumor growth. Performing sensitivity analysis (**Material and Methods**), we identified four model parameters causing a strong variation in the final simulated tumor load (**Fig. 6A**). These sensitive model parameters are: tumor promotion by CAFs (*CFprom*), migration probability of anti-tumor M1-like macrophages (*M1pmig*), tumor mutation load required for macrophage differentiation *(TUthrshM)*, and mutation probability for luminal cells (*TUpmut*). Looking at the dynamics of tumor formation when tuning these parameters, we observed that the mutation probability increases growth speed from the start of the simulation, while the pro-tumorigenic effects of macrophage influx and CAF involvement occur at a later stage **(Supplementary Fig. S3)**. Since these parameters can be related to molecular markers which are largely variable between patients, we decided to vary the corresponding parameters to generate relevant *in silico* patient populations. Analyzing the combined effect of parameter pairs on tumor growth, we empirically selected one high and one low value for each parameter (**Supplementary Table 3**). We chose values for which the effects of the parameter variation were clearly observable, but not too overpowering (other parameters having little/no effect based on **Supplementary Fig. S4**). To reduce the number of variables in order to have big enough clinical patient groups for the analysis described in the next section, we merged the two macrophage parameters: high migration probability and low threshold for phenotype switching (pro-tumor macrophages) versus low migration probability and high threshold for macrophage phenotype switching (anti-tumor macrophages). This resulted in three parameter sets that allow for simulation of patients with: 1. High vs low level of tumor-promoting effect of CAFs; 2. High vs low pro-tumor macrophage characterization; 3. High vs low level of mutation frequency of tumor cells. By systematically combining the effect of these three parameter sets, we obtained eight patient groups (**Fig. 6**).

**Figure 6.**
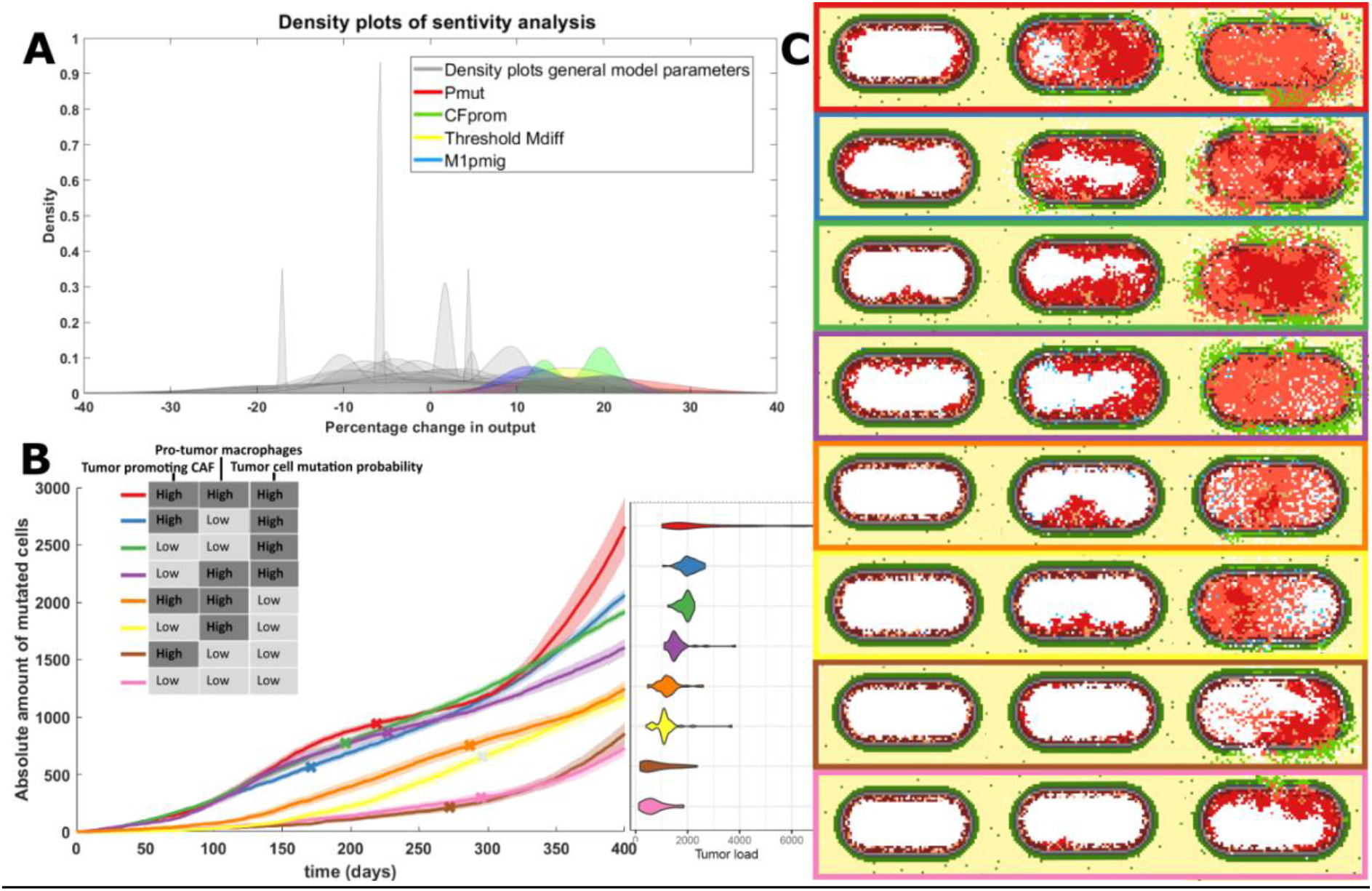
Effect on tumor growth of varying sensitive model parameters. **A**, Grouped histogram of the repeated sensitivity analysis (5 times for each parameter), overlapped by four (differently colored) histograms of the most sensitive parameters: mutation probability of luminal cells (Pmut, red), probability of CAFs promoting tumor cell proliferation (CFprom, green), yellow represents the amount of mutations needed before tumor cells affect macrophage differentiation (TUthrshM) and M1 macrophage migration probability (M1pmig, blue). **B**, The averaged evolution of the amount of tumor cells for 40 simulations that developed cancer for each of the eight subclasses. These classes were based on the ‘high’ or ‘low’ status of sensitive parameters for CAFs, TAMs and tumor cells. Included is a violin plot depicting the spread of simulated tumor cell amounts. **C**, An example of tumor development for each group at an early point in the simulation (50 days), the point at which it becomes invasive and the state at the end of the simulation (400 days).

For all four groups with high tumor mutation probability, over 88% of the simulations showed disease progression towards cancer (**Supplementary Table 4**). This is lower for other groups, with the two groups with pro-tumor macrophages and low mutation probability resulting in modeled cancer progression in less than 8% of the simulations.

To compare model simulations with clinical data, which are only available for developed tumors from patients who underwent prostate surgery, we performed follow-up analysis considering only the simulations resulting in cancer development. The group with the most aggressive tumors consists of simulated patients with high tumor-promoting CAFs, high pro-tumor macrophages characterization and a highly aggressive tumor cell phenotype (red line showing the simulated tumor growth over time in **Fig. 6B** and corresponding example simulation in the red box in **Fig. 6C**). On the contrary, the group with the least aggressive tumors is simulated when all parameter sets are set to ‘low’ (i.e. the least tumor-promoting phenotype; pink line in **Fig 6B** and pink box in **Fig. 6C**).

As expected, the time of invasiveness (i.e., breakdown of the basement membrane, marked with an x in **Fig. 6B**) is significantly earlier for the tumors with high mutation probability as compared to those with low mutation probability (one-sided Wilcoxon Rank Sum test, p-value = 2.26e-10). However, the time of invasiveness does not always correlate with growth speed. The tumor group with the steepest growth curve (red line, **Fig. 6B**) becomes invasive later compared to more slowly growing tumors (e.g., the blue line, with anti-tumor macrophage characterization, p-value = 0.030). This analysis suggests that different mechanisms can affect how quickly tumors develop and how long it takes for tumors to become invasive.

### Model simulations of tumor load associate with patient prognosis

Considering the same eight patient groups (all possible combinations of the three parameter sets) defined in the previous section, we wanted to assess if the *in silico* behaviors correlate with patient prognosis. To do so, we compared model predictions of tumor load (only for cases that developed cancer) with clinical data from a cohort of PCa patients (N=494) from the TCGA database. For each of the three parameter sets we defined whether a patient belonged to the “low” or “high” group considering three molecular markers (see **Supplementary Table 5** for detailed motivation of the choice of the markers). Tumor aggressiveness was defined based on TMB and the expression of two frequently mutated genes in PCa (TP53 and CDKN1B) (17,79,80). Pro-tumor macrophage characterization was defined based on the ratio of M2:M1 macrophages and the expression of two genes involved in pro-tumor macrophage differentiation (CXCL2 and STAT3) (81–83). Finally, the tumor-promoting CAFs effect was defined based on the quantification of CAFs and the expression of two soluble molecules secreted by CAFs that affect tumor progression (TGFBR2 and IGF1; the latter one with an inverse relationship) (50,84–86). For each parameter set, a patient was assigned to the ‘high’ category if at least two out of three makers were above the cohort median, and ‘low’ otherwise. In this way, we could divide the TCGA patients in eight clinical patient groups with similar characteristics to the *in silico* groups.

We observed a negative correlation between the tumor load from the *in silico* patient groups and the PFS time of the matching clinical PCa patients. (Pearson correlation = -0.73, p-value = 0.04, **Fig. 7A**). Patients classified in the three groups with highest tumor load showed a worse prognosis (albeit not statistically significant, p-value=0.089; Kaplan Meier plot in **Fig. 7B**) and a significantly higher Gleason score (chi-squared test, p-value=0.005; **Fig. 7C**) as compared to the patients in the three groups with lowest tumor load. Overall, these results indicate that tumors which are characterized to be more aggressive *in silico* correspond to patients with higher grade and worst prognosis.

**Figure 7.**
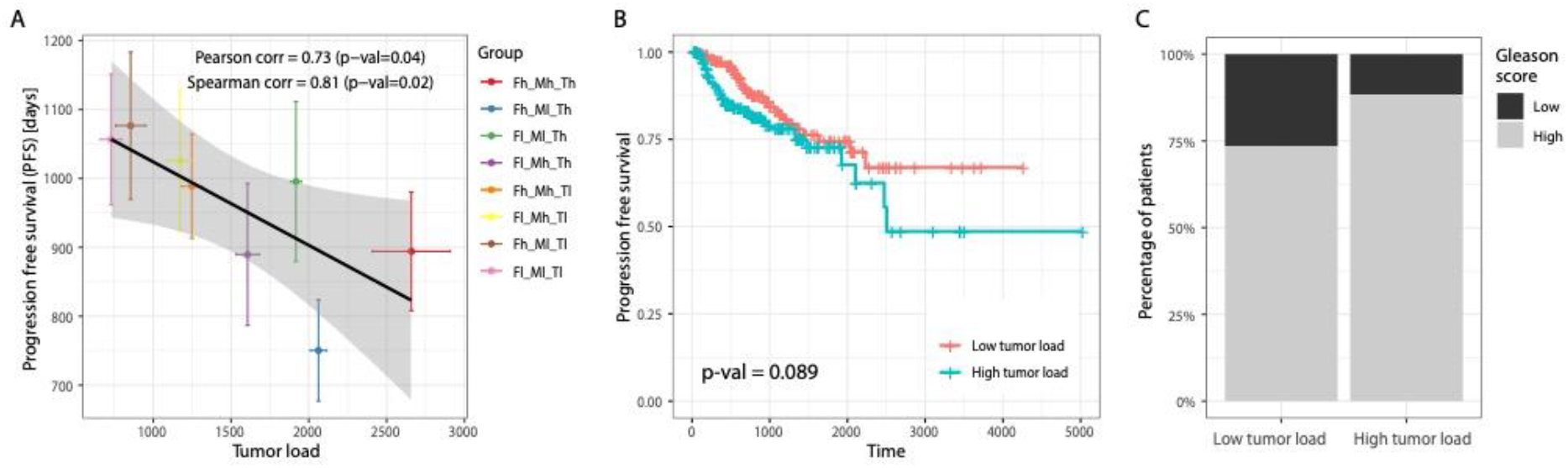
Clinical validation of model predictions for different patient groups. **A**, Correlation between the simulated tumor growth (simulation time 400 days, 40 simulations per modeled patient group) and the average progression free survival time for clinical patients assigned to the matching patients groups based on molecular markers. Colors correspond to those used in Figure 6B, portraying simulated tumor growth over time of the same classes. **B**, Kaplan Meier plot of two patient groups. Patients were considered as low tumor (red) load if they belong to the three groups with lowest simulated tumor load and high tumor load (blue) if they belong to the three groups with highest simulated tumor load. **C**, Binary Gleason scores per patient group; Gleason scores of 7 or higher were considered ‘high’ and Gleason scores of 6 or lower were considered ‘low’.

## Discussion

The process of PCa development can take years and is heavily influenced by many different types of cells, stochastic events, and the tumor microenvironment. Its unpredictable nature and extensive adaptation strategies bear resemblance to the process of evolution, which makes it particularly hard to combat at a later stage. Recreating the complete disease settings to better understand and treat the disease is therefore rather difficult in *in vitro* or *in vivo* settings.

As recently emphasized in an opinion paper by West and colleagues (87), agent-based models are key tools to reproduce the complexity of the tumor *in silico*, offering a complementary approach to *in vitro* and *in vivo* experiments. They allow the integration of different types of knowledge, framing it in the form of an intuitive set of rules. Despite their simplicity in the formulation, they allow simulation of complex behaviors deriving from cell-cell interactions.

Here, we designed a comprehensive agent-based model that provides an *in silico* experimental set up to study PCa onset and progression. The rules defining our ABM were based on a set of assumptions integrating knowledge from several studies. Model parameters were additionally fine-tuned by fitting our in-house generated *in vitro* co-culture data. After showing that our model was able to reproduce known tumor patterns and relevant steps of tumor progression, we used the model to *in silico* study the impact that deterministic and stochastic events have on PCa progression.

In our study we identified pro-tumor activity of CAFs and macrophages and mutation probability of the tumors as main deterministic causes of *in silico* tumor heterogeneity. While high tumor mutation probability generally results in fast invasion and bigger tumors, the effects and quantities of macrophages and fibroblasts at different time points were found to be a very important factor in PCa development and progression too. These findings could help to improve our understanding of different patient molecular characteristics and how these contribute to the likelihood of progression, thus suggesting new prevention strategies and options for patient-tailored treatment plans. However, more clinical data on patients not (yet) in a malignant disease stage would be needed to assess if these markers could be used as indicators of disease stages and be functionally associated with disease progression. This assessment could be tested by monitoring prostatitis patients, which is a risk factor for PCa (88).

We additionally observed that, running the model multiple times starting with the same initial conditions, only a fraction of the simulations developed into cancer. This is determined only by the stochasticity of the events included in the simulation that mimics the *in vivo* stochasticity of cellular interactions. We observed that aleatory events related to the interactions between macrophages and tumor cells can determine the success of early immunosurveillance thus determining the fate of the tumor. The stochasticity of interactions also affects how long it takes before the tumor becomes invasive, driven by the balance between the number of CAFs, amount of driver mutations and the ratio of anti-/pro-tumor macrophages. While there is increasing awareness that clinicians should consider the impact of genetics to account for patients heterogeneity in prostate cancer management (89,90), our results underlie the importance of monitoring the microenvironment phenotype (e.g. using multiplexed tissue imaging) during PCa progression.

Although we have shown that our AMB model is a valuable tool to conduct *in silico* experiments on the onset of prostate cancer, it is important to keep in mind that models are always an approximation of reality and the choice of the level of details included is driven by the aim of the study. Our model could be extended in the future to study treatment response and more advanced disease stages, such as the effect of androgen deprivation therapy or androgen receptor (AR) inhibition and the development of castration resistance. Considering that AR is known to play a role, not only on prostate cancer cells, but also on fibroblasts (26) and macrophages (55), an extension of our ABM could be a valuable tool to take an integrative approach to study how the the PCa microenvironment mediates therapy response.

Additionally, for this study we chose to focus on macrophages and fibroblasts because of their prominent role in PCa, but the model could be further extended to include other cell types, such as T-cells. Although PCa is known to be an immune excluded and suppressed tumor type, recent studies showed the potential of combining T-cell-based immunotherapies (i.e. immune checkpoint blockers or CAR T cells) with other therapies targeting the PCa microenvironment to restore anti-tumor immunity in advanced prostate cancer (91,92). ABMs could help to understand the effect of combining different therapies in specific microenvironment subtypes, therefore suggesting how to tailor combinatorial treatment.

Furthermore, we have now chosen to model the effect of cytokines and chemokines implicitly (e.g. by basing an interaction between two cells on the distance between them), but it would be an interesting addition to model humoral factors explicitly (e.g. using hybrid models (93)), for example when wanting to zoom in more on androgen dependence and the path to castration resistant disease. However, this would also increase the number of model parameters and the computational costs.

Previous *in silico* models of PCa have been focused on specific mechanisms such as the formation of bone metastases (76) or the role of disrupted stem cell movement in causing excessive growth in healthy prostatic ducts (94). To our knowledge, this is the first ABM to simulate the onset and development of prostate cancer in healthy prostatic acini considering the effects of the microenvironment including fibroblasts and macrophages. Our analysis shows that, not only tumor cells, but also macrophages and fibroblasts play an important role in PCa development and could provide potential markers of disease progression.

## Acknowledgements

The results shown here are in part based upon data generated by the TCGA Research Network: http://cancergenome.nih.gov/. The authors would like to thank Oscar Lapuente-Santana for providing the pre-processed TCGA data, dr. F. Finotello for assistance in running the *immunedeconv* package and J. van Leeuwen for testing the code of the ABM. We would like to thank members of the Eduati, Zwart and Bergman labs for valuable discussion and feedback.

