## Supplementary Tables S1-S5 and Supplementary Figures S1-S3 for "Exploring the onset and progression of prostate cancer through a multicellular agent-based model"

Author affiliations

*Supplementary Table S1. List of model assumptions.*

| **Model assumption** | **Reference** |
|---|---|
| The shape and ratios of cells in the starting geometry; a prostatic acinus | (1) |
| The prostatic acinus is surrounded by ECM | (2) |
| ECM has variable permeability, depending on the stage of the disease | (2) |
| The presence and percentage of prostate cancer stem cells | (3,4) |
| Tissue resident fibroblasts are quiescent and become active upon receiving a signal | (5) |
| Basement membrane is quiescent and functions as a barrier between the inside and outside of the acinus. | (1,6) |
| Quiescent cells remain in equilibrium/homeostasis; they do not migrate, proliferate or die | Trivial |
| Basement membrane is broken down by proteolytic enzymes produced by tumor cells. Not all enzyme production results in actual break down (only 10-30 percent) | (6) |
| Basement membrane break down does not happen right away, so tumor cells have to gain a certain amount of mutations before they can start break down | (6) |
| Basal cells and non-mutated luminal cells are in homeostasis; they can die and proliferate at low rates, within their normal/healthy regions | Trivial |
| Basal cells should always be attached to the basement membrane | (6) |
| Basal cells can only die after enough luminal cells have mutated | (7) |
| Non-quiescent cells can spontaneously enter apoptosis and die | (8), Experiments |
| Luminal cells can gain mutations | (1) |
| Mutated luminal cells have a higher chance of gaining another mutation and are not quiescent or in equilibrium | (1) |
| Luminal stem cells can also gain mutations, generating ’cancer stem cells’ | (1) |
| Mutated cells can proliferate | Experiments |
| Upon gaining a mutation, oncogenic or tumor suppressing genes are affected. Mutated cells gain a proliferative advantage | (9) |
| Apart from the ’standard’ actions proliferate,migrate and die, mutated cells can perform additional actions (parallel to the other actions) once they have gained enough mutations | Trivial |
| Additional actions of mutated cells include: affecting differentiation of macrophages, breaking down basement membrane and affecting ECM permeability | (2,6,10) |

| | |
|---|---|
| Mutated cells can only break down basement membrane if they are directly adjacent to it | Assumption |
| Mutated cells move by a random walk and migrate towards fibroblasts | Experiments, (11) |
| Activation of tissue resident fibroblasts happens during PIN and is affected by mutated cells. CAFs co-evolve during disease progression. | (12–15) |
| The effect of CAFs on tumor cells depends on the distance between the two | (16) |
| CAFs can migrate towards mutated cells | (8) |
| CAFs can proliferate and migrate more than normal fibroblasts | (5) |
| CAFs stimulate primary tumor growth | (17) |
| CAFs elicit Epithelial Mesenchymal Transition (EMT) in mutated cells | (18) |
| CAFs modify the ECM; making it rather impermeable at the beginning and produce MMPs later on to degrade it | (2,13,19) |
| Macrophages cannot proliferate (the few times they possibly could does not affect simulations) | Experiments |
| M1 macrophages enter the simulation from one point, mimicking entry via a blood vessel | (20) |
| Macrophages migrate towards mutated cells, but only if they are in reach (17 grid spaces). If they are further away, they move randomly | (8,19) |
| Macrophages infiltrate prostatic acini during PIN | (21,22) |
| Macrophages can kill mutated cells. M1 polarized macrophages are more likely to kill than M2 polarized macrophages | Experiments, (23) |
| If a macrophage kills a mutated cell, more macrophages are drawn to the acinus (increased influx) | (21,22) |
| M2 macrophages do not enter the simulation via the blood, but have to arise from differentiation of the M1 macrophages | (20) |
| M2 polarized macrophages promote mutated cell proliferation | Experiments, (20,21) |
| Macrophages have to be adjacent to mutated cells to be able to promote those or kill those | (21,24) |
| M2 macrophages can elicit anchorage independent growth in tumor cells | (21,25) |

*Supplementary Table S2. List of all model parameters.*

| Model parameter | Description | Value | Reference |
|---|---|---|---|
| LCpprol | Proliferation probability for luminal cells | 0.3054 | Experiments, A |
| TUpprol | Proliferation probability for mutated luminal cells | 0.3054 | Experiments, A |
| TUpmig | Migration probability for luminal cells that went through EMT | 0.3501 | Experiments, (8) |
| LCpdeath | Probability of death for luminal cells | 0.00284 | Experiments, A |
| TUpdeath | Probability of death for mutated luminal cells | 0.00284 | Experiments, A |
| TUrwalk | Random walk effect on movement | 0.5 | Experiments, (8) |
| TUpmax | Maximum amount of times luminal cells can proliferate (without mutations) | 4 | Experiments, A |
| TUps | Probability of symmetric division if proliferation for stem cells | 0.2967 | B |
| TUpmut | Probability of gaining a mutation for luminal cells | 0.00015 | (26), B |
| TUantig | Added antigenicity per mutation | 0.005 | B |
| TUmutmax | Maximum amount of antigenicity luminal cells can obtain | 1 | B |
| TUadded values | Added probability for gaining a mutation (for cells that have one or more mutations already) | 0.15*TUa ntig | (26) |
| TUadded values | Added maximum proliferation capacity (eac time an additional mutation is gained) | 0.75 | B |
| TUadded values | Added proliferation probability | 0.001* TUantig | B |
| TUpkill | Probability of breaking down the basement membrane by mutated cells | 0.3 | (6), B |
| TUthrshBM | Needed amount of mutations before basement membrane can be broken down | 6 | Arbitrary, C |
| TUpmdiff | Probability of affecting macrophage differentiation by mutated luminal cells | 0.8 | (10,27) |
| TUthrshM | Needed amount of mutations before macrophage differentiation can be affected by mutated cells | 14 | Arbitrary, C |
| TUpMMP | Probability of breaking down surrounding ECM | 0 | Variable, D |

| | | | |
|---|---|---|---|
| TUthrshMMP | Needed amount of mutations before mutated cells can affect ECM break down | 15 | Arbitrary, C |
| M1kmax | Maximum killing capacity of M1 macrophages | 11 | Experiments, A |
| M1pkill | Probability of killing by M1 macrophages | 0.0918 | Experiments, A |
| M1pmig | Migration probability of M1 macrophages | 0.8001 | (26) |
| M1pdeath | Probability of death for M1 macrophages | 0.0147 | Experiments, A |
| M1rwalk | Random walk effect on movement of M1 macrophages (if further away than 17 gridspaces) | 0.8 | (21), B |
| M1speed | Speed of movement | 40 | (26), B |
| M1engagement Duration | Number of steps the M1 cell is engaged | 60 | (26), B |
| M1influxRate | Amount of macrophages that enter the simulation each time | 1 | (8) |
| M1influxProb | Probability of M1 macrophages entering the simulation | 0.08 | B |
| M1influxadd | Factor with which the influx increases after killing the first mutated cell | 3 | B |
| M2kmax | Maximum killing capacity of M2 macrophages | 11 | Experiments, A |
| M2pkill | Probability of killing by M2 macrophages | 0.0381 | Experiments, A |
| M2pmig | Migration probability of M2 macrophages | 0.8001 | (26) |
| M2pdeath | Probability of death for M2 macrophages | 0.0147 | Experiments, A |
| M2rwalk | Random walk effect on movement of M2 macrophages (if further away than 17 gridspaces) | 0.8 | (21), B |
| M2speed | Speed of movement | 40 | (26), B |
| M2influxProb | Probability of M2 macrophages entering the simulation | 0 | (27) |
| M2TUadd | Added probability for proliferation of luminal cells | 0.2985 | Experiments, A |
| M2engagement duration | Number of steps the M2 cell is engaged | 60 | (26), B |
| M2emt | Probability of the M2 macrophage eliciting EMT in a mutated luminal cell if in range | 0.75 | (25), B |
| Fpprol | Proliferation probability of fibroblasts | 0 | (5), E |
| Fpmig | Migration probability of fibroblasts | 0 | (5), E |
| Fpdeath | Probability of death of fibroblasts | 0 | (5), E |

| | | | |
|---|---|---|---|
| Fpmax | Maximum proliferation capacity | 0 | (5), E |
| Frwalk | Random walk effect on movement of fibroblasts | 0 | (5), E |
| Fdiff | Probability of fibroblasts turning into CAFs when in range of mutated luminal cells | 0.75 | (28) |
| CFpprol | Proliferation probability of activated fibroblasts (CAFs) | 0.00838 | Experiments, B |
| CFpmig | Migration probability of CAFs | 0.2 | B |
| CFpdeath | Probability of death of CAFs | 0.0054 | (5), Experiments |
| CFpmax | Maximum proliferation capacity | 4 | Experiments, A |
| CFrwalk | Random walk effect on movement of CAFs | 0.5 | Experiments, B |
| CFemt | Probability of the CAF eliciting EMT in a mutated luminal cell if in range | 0.75 | (18),B |
| CFprom | Added probability for proliferation of mutated luminal cells | 0.5* TUpprol | (17) |
| CFmmp | Probability of surrounding ECM break down by CAFs | 0.9 | B |
| CFmdiff | Probability of CAFs affecting macrophage differentiation | 0.5 | Variable, D |
| Cpprol | Proliferation probability of basal cells | 0.2 | (7), F |
| Cpmig | Migration probability of basal cells | 0 | (7), F |
| Cpdeath | Probability of death of basal cells | 0.01 | (7), F |
| Cpmax | Maximum proliferation capacity | 10 | B |
| Bpdeath | Probability of death of basement membrane | 0 | Trivial |
| Bpprol | Probability of proliferation for basement membrane | 0 | Trivial |

***A*** - These parameters were determined by fitting them to the obtained experimental data, using the PSO algorithm described in the Materials and Methods section.
***B*** - These parameters can be freely chosen by the user. They were qualitatively tuned for these simulations to yield realistic results.
***C*** - These parameters (thresholds) can be freely chosen by the user. Increasing these parameters increases the amount of time it takes before disease can progress towards the next stage.
***D*** - These parameters were set to zero for the performed simulations, but can be set to higher values if the user wishes to include these processes/effects.
***E*** - These fibroblast parameters were all set to zero, as the tissue resident fibroblasts are believed to be quiescent before they are activated as CAFs in this case (5).

***F*** - Disruption of the basal cell layer takes place exponentially; only 0.7 percent breakdown in the first stage of PIN, 15 percent in the second stage and 52 percent in the third stage. These parameters were qualitatively tuned to get values close to these.
***G*** - Distances from which mutated cells affect macrophage differentiation and elicit ECM break down, the distances from which CAFs can break down ECM or elicit EMT and the distance from which macrophages can elicit anchorage independent growth were found to be insensitive during the sensitivity analysis. The chosen values are trivial (if bigger than 1).

*Supplementary Table S3. Parameter values for the eight patient phenotypes.*

| **Parameter** | **Low/anti tumor value** | **High/pro tumor value** |
|---|---|---|
| TUpmut | 0.0000075 | 0.000175 |
| CFprom | 0.01 | 0.7 |
| M1pmig<br>TUthrshM | 0.01<br>20 | 0.51<br>1 |

*Supplementary Table S4. Percentage of the simulations that results in cancer for all eight phenotypes.*

| **Simulated patient phenotype** | **Percentage of cancer development** |
|---|---|
| High CFprom, pro-tumor macrophages & high Pmut | 89 % |
| High CFprom, anti-tumor macrophages & high Pmut | 96% |
| Low CFprom, anti-tumor macrophages & high Pmut | 96% |
| Low CFprom, pro-tumor macrophages & high Pmut | 89% |
| High CFprom, pro-tumor macrophages & low Pmut | 7.5% |
| Low CFprom, pro-tumor macrophages & low Pmut | 7.9% |
| High CFprom, anti-tumor macrophages & low Pmut | 27% |
| Low CFprom, anti-tumor macrophages & low Pmut | 23% |

*Supplementary Table S5. Patient markers distinguishing the eight different patient classes*

| Marker | Related model parameter | Description | References |
|---|---|---|---|
| Macrophage-related markers: below, we describe the patient markers that we chose to determine whether a patient would either have more tumor promoting macrophages (high M2/M1 fraction and high levels of CXCL2 and STAT3) or less tumor promoting macrophages (low patient marker levels). | | | |
| Fraction M2/M1 | *M1pmig* *TUthrshM* | This fraction describes the ratio of tumor promoting macrophages (M2) to the amount of anti-tumor macrophages (M1). It shows the proportion of macrophages that transitioned towards the tumor-promoting phenotype. | |
| CXCL2 | *M1pmig* *TUthrshM* | CXCL2 is a chemokine involved in the differentiation process of macrophages towards their pro-tumor phenotype. Indicating whether (many) macrophages are transforming towards the pro-tumor phenotype. | (29,30) |
| STAT3 | *M1pmig* *TUthrshM* | STAT 3 is a transcription factor involved in the differentiation process of macrophages towards their pro-tumor phenotype. Indicating whether (many) macrophages are transforming towards the pro-tumor phenotype. | (31) |
| Tumor cell related markers: below, we describe the patient markers that relate to the mutation probability. On the basis of these markers, we determined whether a patient had high mutation probability (high mutational burden and high TP53 and CDKN1B) or low mutation probability (low patient marker levels). | | | |
| Mutational burden | *TUpmut* | An indication of the amount of mutations that have occurred in the tumor cells (this thus is an indication of the frequency with which the patients' cells mutate). | |
| TP53 | *TUpmut* | TP53 is a gene that is commonly mutated in PCa. Especially mutations in exon 7 and 8 contribute greatly to disease progression and recurrence. | (9,32) |
| CDKN1B | *TUpmut* | CDKN1B is a kinase inhibitor that correlates with tumor progression and PSA levels. It is a tumor suppressor gene; loss of heterozygosity has been detected in approximately 50% of prostate tumors. | (33,34) |
| CAF related markers: below, we describe the patient markers that relate to the tumor promoting ability of CAFs. On the basis of these markers we determined whether a patient had more tumor promoting CAFs (high fraction of CAFs, high TGFBR2 and low | | | |

| | | | |
|---|---|---|---|
| IGF1) or less tumor promoting CAFs (low fraction of CAFs and TGFBR2 and high IGF1). | | | |
| Fraction of CAFs | *CFprom* | This fraction describes the number of CAFs present. | |
| IGF1 | *CFprom* | IGF1 is a growth factor secreted by CAFs that plays a role in tumor progression. An inverse relation was found between (the presence of) PCa and levels of IGF1. | (35,36) |
| TGFBR2 | *CFprom* | TGFBR2 is involved in the activation of resident fibroblasts towards CAFs and allows for crosstalk of the tumor microenvironment cells. | (14,37,38) |

*Patient markers that were used to create the eight different groups of patients. For each class (the first three rows are class: 'TAM', row 4-6 are class 'tumor cells' and row 7-9 are 'CAFs') the three markers were compared to the median value for all included patients. If two (or more) out of three markers were below median value, the patient was marked as 'low' for that class (e.g. low TAM markers means low in terms of TAM, which equals less tumor promoting macrophages). If two (or more) out of three markers were above median value, the patient was marked as 'high' for that class. Combining all the possible combinations (Ranging from all classes high to all classes low) yields the eight different patient phenotypes that were also simulated using the In Silico model. The model parameters that these classes relate to can be found in table S2. M1pmig indicates migration probability of M1 macrophages, TUthrshM indicates the required amount of mutations before a tumor can affect macrophage migration, TUpmut indicates the mutation probability of tumor cells and CFprom indicates the tumor promoting probability by CAFs.*

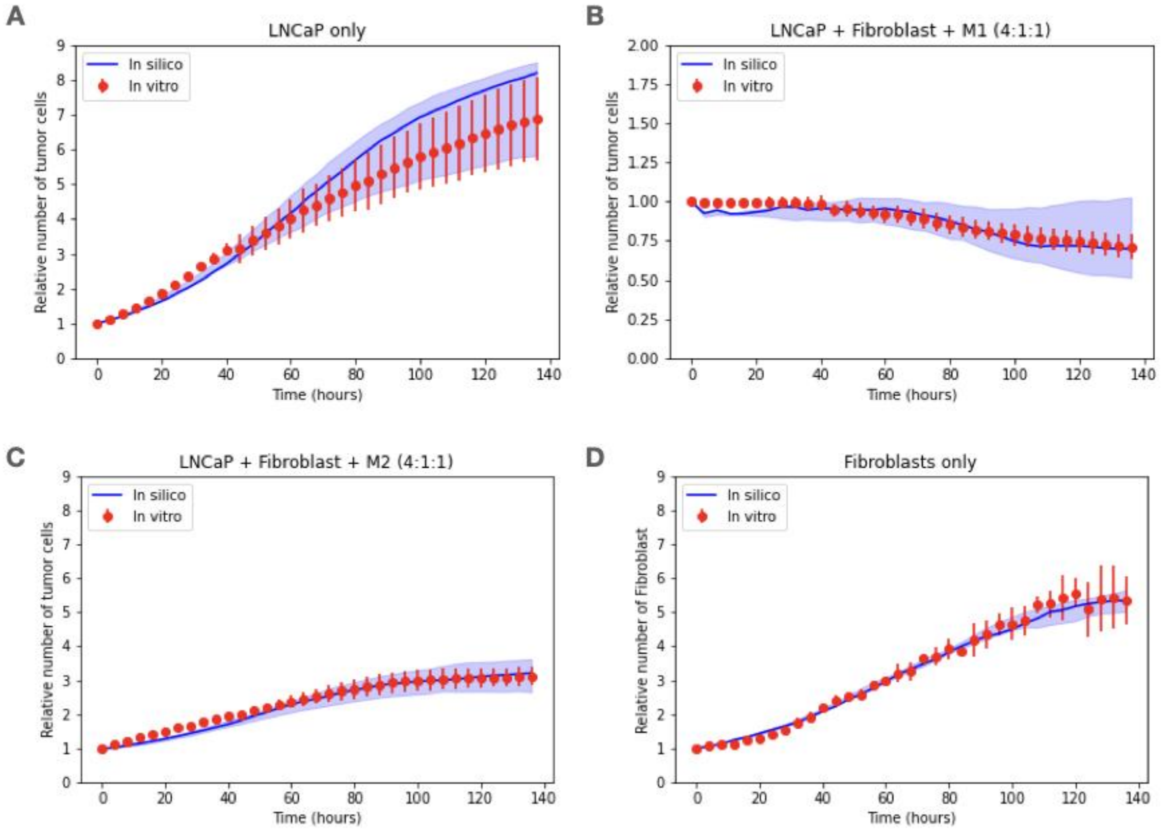


*Supplementary Fig. S1. Experimental data and model fitting. Growth curves of A. LNCaP cells; B. LNCaP co-cultured with fibroblasts and M1 macrophages; C. LNCaP co-cultured with fibroblasts and M1 macrophages; D. Fibroblasts. Red dots represent means and error bars represent standard deviation of the three biological replicates (spanning six technical replicates). Blue lines represent model simulations using the median of the parameters estimated for the biological replicates, and shading represents the interquartile range for the 50 optimisations per biological replicates.*

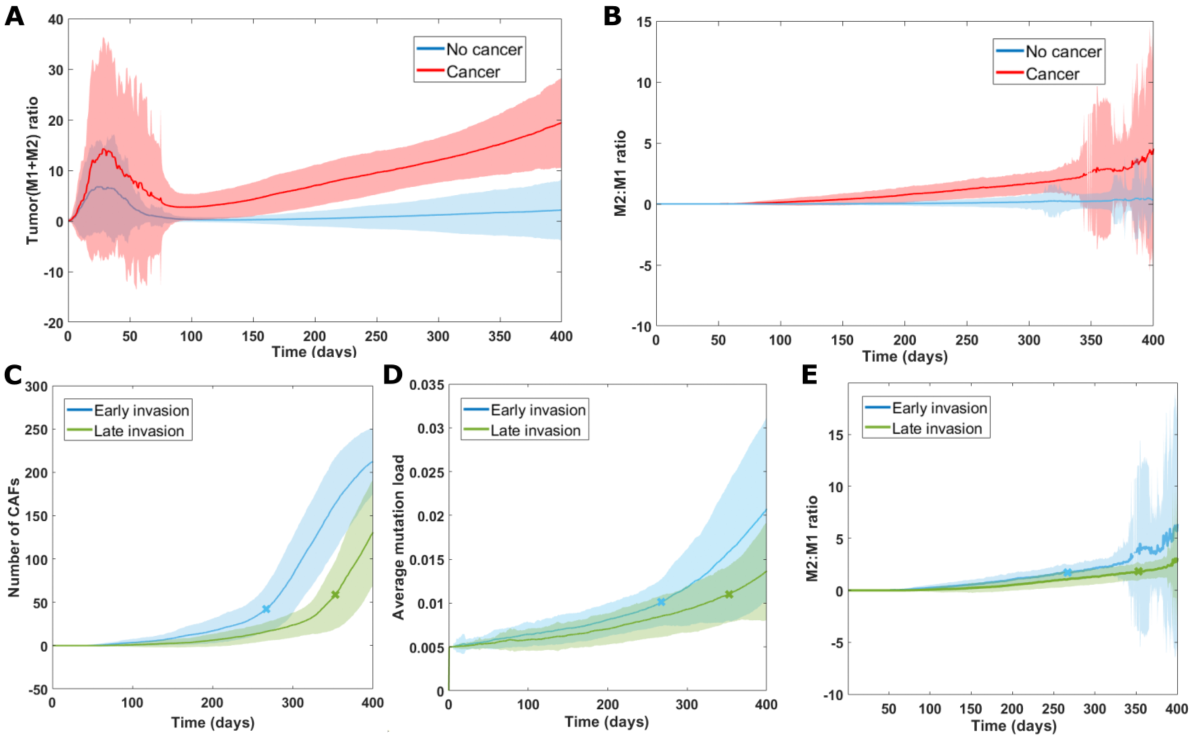


*Supplementary Fig. S2. Cancer and non cancer simulation comparison. Average results (including standard deviation in the shaded area) for simulations using the standard parameter values (as described in table S2). A. Tumor cell to macrophage ratio (M1+M2) both for the simulations that resulted in cancer (180/500) and that did not result in cancer (320/500). B. Ratio of M2:M1 macrophages for non-cancer and cancer cases. C. Number of CAFs over time for all cancer cases in groups of early and late invasion (invasion time indicated with a cross). D. Average mutational load for all cancer cases of early and late invasion (invasion time indicated with a cross). E. M2:M1 ratio for all cancer cases (invasion time indicated with a cross).*

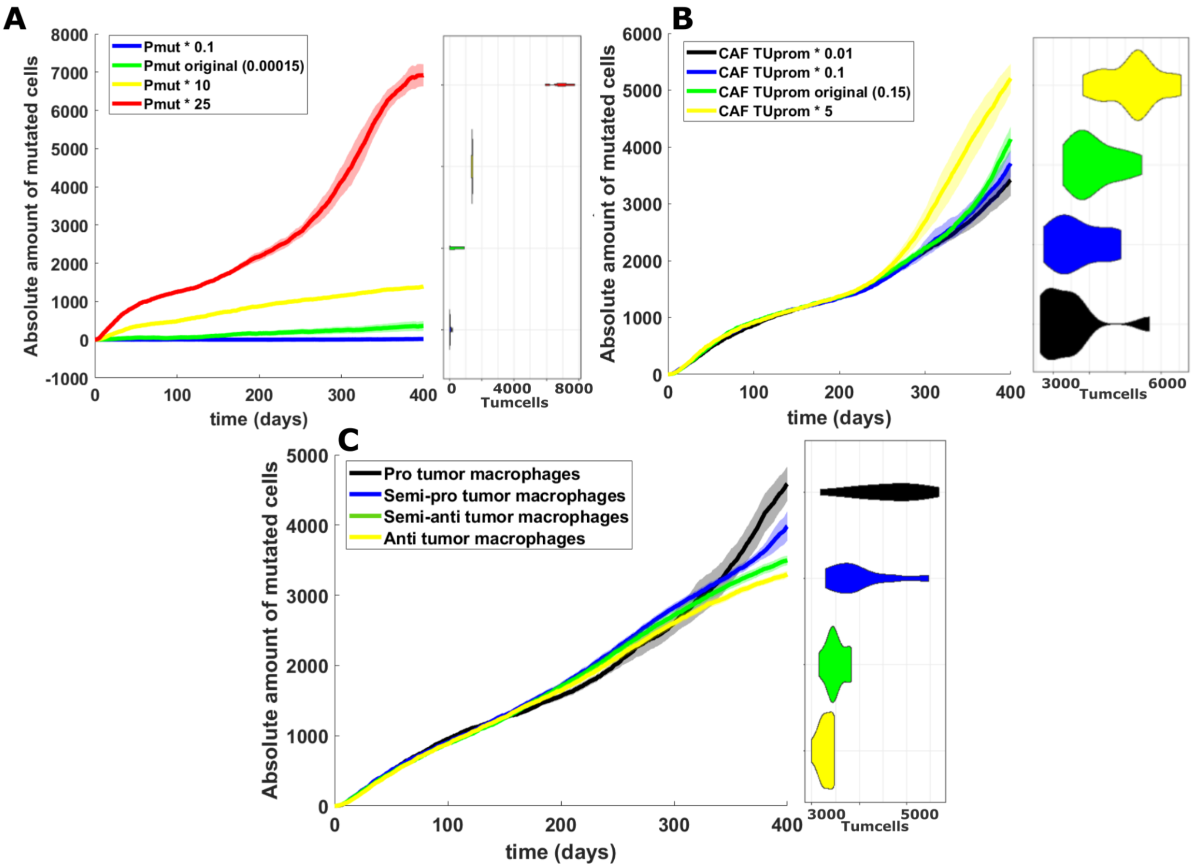


*Supplementary Fig. S3. Follow-up sensitivity analysis. A. Sensitivity analysis of the probability to gain mutations for luminal cells. Next to this graph, the violin plot on the four different simulations is displayed. B. Sensitivity analysis of the probability for mutated cell proliferation promotion by CAFs. The violin plot is displayed next to the graph, the colors correspond to those displayed in the graph. C. Sensitivity analysis of the combined parameters for macrophages (M1 migration probability and mutation threshold). Pro-tumor macrophages have a lower threshold for transition towards pro-tumor phenotype and larger migration probability. Next to the graph, the violin plot for the four simulations is displayed.*

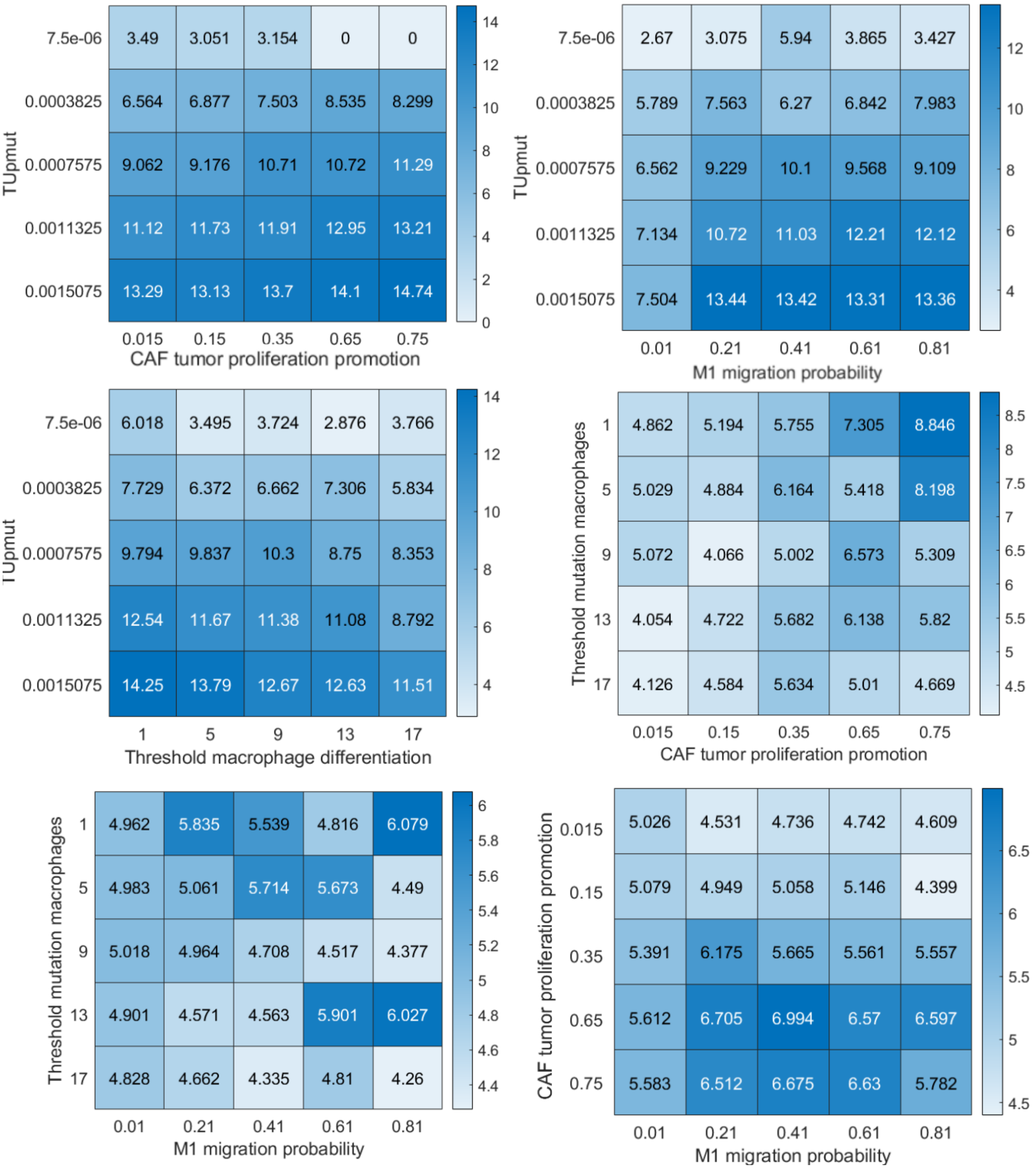


*Supplementary Fig. S4. Pairwise combinations. Heatmaps of the pairwise combinations of selected model parameters. The darker the square (and higher the number), the larger the tumor.*

30. Nagarsheth N, Wicha MS, Zou W. Chemokines in the cancer microenvironment and their

relevance in cancer immunotherapy. Nat Rev Immunol. 2017;17:559–72.
